# The genetically-encoded amino acids distribute non-randomly within a functionally-relevant chemical space

**DOI:** 10.64898/2026.05.06.723277

**Authors:** Sean M. Brown, Judson Hervey, Scott N. Dean, Gary J. Vora

## Abstract

The standard set of 20 genetically-encoded amino acids (C20) exhibits a statistically non-random distribution in primarily two structurally-relevant physicochemical properties: hydrophobicity and molecular volume, and to a lesser extent charge. It remains an open question, however, whether evolutionary pressures similarly optimized the same alphabet for the distribution of functionally-relevant properties, such as reactivity. In this study, we used semi-empirical quantum chemistry simulations to calculate the highest occupied molecular orbital and lowest unoccupied molecular orbital (HOMO-LUMO) gaps for 84 xeno amino acids and constructed 10 million random 20-mer amino acid alphabets to determine where C20 fit amongst this background. The HOMO-LUMO gap measurements demonstrated that C20, similar to hydrophobicity and volume, also exhibits a non-random distribution. However, unlike hydrophobicity and volume, this distribution is non-random across an unevenly broad range. The results expand upon previous theory and suggest HOMO-LUMO gap energies as one synthetic biologists may consider when developing novel protein design tools or designing functional xeno amino acid alphabets.

**Highlights:**

- Life’s amino acid alphabet is non-randomly distributed within an expanded computationally-generated chemistry space generated from large-scale quantum chemistry simulations.
- Amino acid alphabet coverage theory applies beyond structurally-relevant physicochemical descriptors to include functionally-relevant properties like reactivity as measured by frontier molecular orbitals
- Findings here provide a theoretical framework to guide the design of novel proteins and development of synthetic amino acid alphabets.

## Introduction

The 20 genetically-encoded amino acids (C20) exhibit a statistically non-random distribution, or coverage, in van der Waals’ volume (*i.e*. size), logP (*i.e*. hydrophobicity), and to a lesser extent charge.^1– 4^ Coverage is defined by the combination of (i) the measured property range for a set of amino acids (AAs), and (ii) the evenness with which they distribute across each range (Fig. 1A). Range allows for chemical diversity, while evenness minimizes the phenotypic jump as one AA substitutes for another in any given sequence, three-dimensional structure or phenotype. Recently, coverage theory was even applied to successfully design, synthesize, and characterize xeno amino acid (xAA) sets with familiar secondary structures such as α-helices.^4^

**Figure 1.**
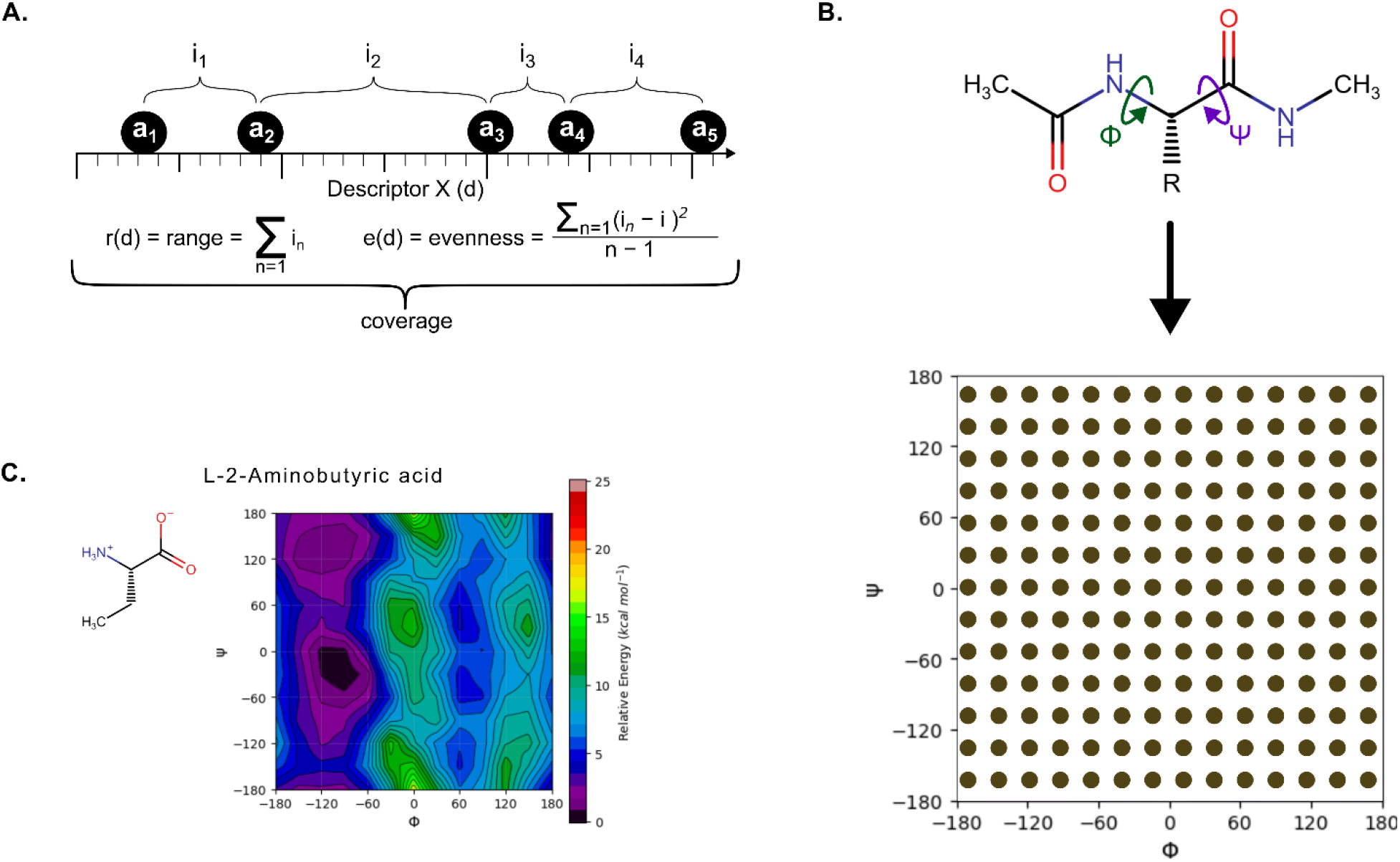
Physicochemical coverage to potential energy surfaces. **A.** Definition of range and evenness. For a given chemical descriptor, such as van der Waals volume, “coverage” joins two statistics to characterize a set of AAs. These statistics are shown for an example set of five AAs (*a*_*1*_*…a*_*5*_) with four corresponding intervals (*i*_*1*_*…i*_*4*_) measured in terms of the hypothetical quantitative ‘Descriptor X’ (*d*). Evenness (*e*) is the sample variance of the intervals between AAs (*i*); Range (*r*) is the sum of these intervals (∑*i*_*1…4*_); “Coverage” is therefore the combination of range AND evenness for any given physicochemical descriptor. **B**. As AAs are covalently linked via peptide bonds, two dihedral angles, phi (Φ) and psi (Ψ) are formed. As starting conformers for PES generation, we iterated through and generated initial geometries for 144 evenly distributed phi and psi angles. **C**. An example of a PES generated for the xAA L-2-Aminobutyric acid. The corresponding relative ECOSMOS DFT single point energy is represented by a linear color gradient from black (0 kcal/mol) to red (25 kcal/mol).

The physicochemical properties of hydrophobicity and volume are critical for the formation of protein structure. Hydrophobic collapse is widely accepted as an underlying principle,^5–7^ even the driver,^8^ of protein folding, whereas volume determines the steric constraints for structure formation. While the three-dimensional structure supports the functional capacity of proteins, relatively few AAs are ultimately responsible for catalysis, or functionality, within a given protein.^9,10^ As such, it is somewhat surprising that the coverage of functionally-relevant physicochemical properties have yet to be investigated to the same extent as structurally-relevant properties (*i.e*. hydrophobicity, volume, and charge).

Within any molecule, the gap in energy between the highest occupied molecular orbital (HOMO) and the lowest unoccupied molecular orbital (LUMO) often is regarded as a measure for chemical reactivity.^11–13^ In this study, we calculate the functionally-relevant property of HOMO-LUMO gaps (E_g_) for many xAAs, *a priori*, using state-of-the-art semi-empirical quantum chemistry simulations. With these values, we then calculated the coverage of 10 million xAA alphabets to probe the statistical randomness of reactivity within C20. The findings suggest that, similar to distributions for hydrophobicity and volume, reactivity is also non-randomly selected for in C20.

## Results

We sought to determine to what extent C20 has been optimized for reactivity as measured by frontier molecular orbital energies (HOMO-LUMO gaps, E_g_). As a first approach, we conducted a Monte Carlo simulation (10 million iterations) comparing C20 to randomly generated 20-member alphabets drawn from a library of 84 xAAs. This library contained xAAs returned from a heuristic search for alphabets comprising AAs predisposed to having hydrophobicity and volume distributions similar to C20 (see Methods). As a control, we repeated previous analyses that investigated the distribution and range of hydrophobicity (JChem logD at pH 7.0) and van der Waals volume (vdW Volume)^1–3,14^. As expected, C20 was not a notable outlier in distribution of hydrophobicity or volume against this particular background. C20 instead occupied a relatively narrow and uneven distribution of volume (12^th^ percentile range, 78^th^ percentile evenness) and a similarly unremarkable narrow and even distribution of hydrophobicity (38^th^ percentile range, 43^rd^ percentile evenness) when compared to the random alternative background (Fig. 2 and Table 1). These data verified that the library of random AA sets was a representative background for viable alternative AAs in terms of a structurally-relevant chemical space (*i.e*. hydrophobicity and volume).

**Table 1.** C20 distribution across descriptors.

| Property | Descriptor | Range Percentile | Evenness Percentile | KS Percentile |
| --- | --- | --- | --- | --- |
| Reactivity | HOMO-LUMO (eV) | 89 | 5 | 0 |
| Volume | vdW (Å<sup>3</sup>) | 12 | 78 | 71 |
| Hydrophobicity | JChem LogD | 38 | 43 | 91 |

vdW = van der Waals volume; JChem LogD = hydrophobicity; KS = Kolmogorov-Smirnov statistic

**Figure 2.**
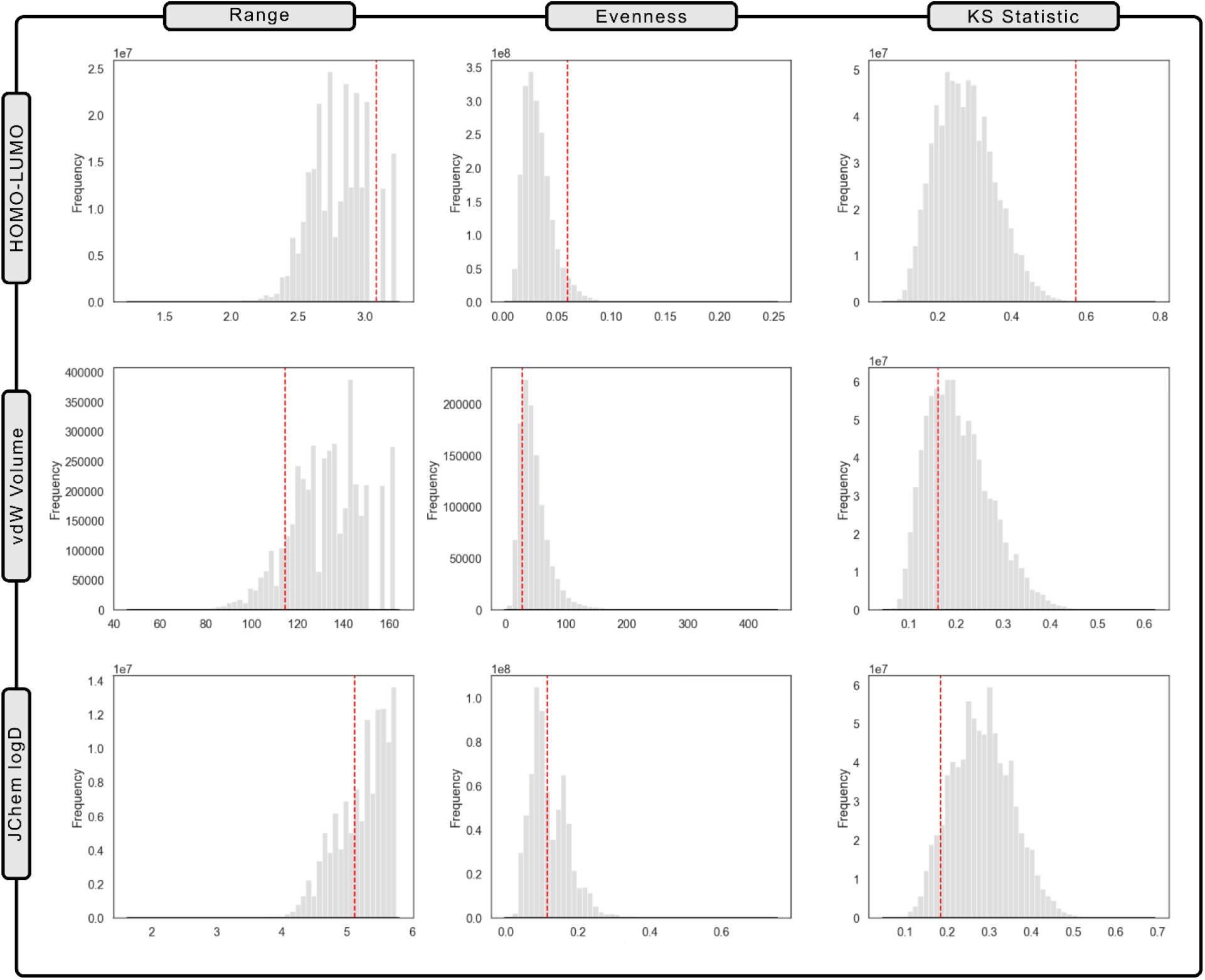
Statistical analysis of C20 physicochemical properties amongst a background of alternatives. Distribution metrics for 10 million randomly generated xeno alphabets of equivalent size (grey histograms) compared to C20 (red dashed line) for various coverage values [Range (left column), Evenness (center column), KS statistic (right column)] and HOMO-LUMO gaps (top row), van der Waals volume (center row) and JChem logD hydrophobicity (bottom row). Percentile values for C20 can be found in Tables 1 and 2.

After confirming this baseline for structurally-relevant properties, we focused on chemical reactivity as measured using HOMO-LUMO gaps. Consistent with previous findings,^15–17^ we observed that the majority of C20 AAs (*e.g*. AGLNKSQTRVIE) have relatively similar E_g_ to one another (Fig. 3A). Exceptions primarily came from AAs with aromatic side chains such as tryptophan (W), histidine (H), and tyrosine (Y). In stark contrast to hydrophobicity and volume, even among alphabets comprising xAAs predisposed to forming C20-like sets, C20 exhibits a non-random range of E_g_ (89^th^ percentile) when compared to the xeno alphabet background (Fig. 2 and Table 2). Unlike C20’s non-random distribution in hydrophobicity and volume, however, the wide range exists within a remarkably uneven distribution (5^th^ percentile in evenness and 0^th^ percentile in the Kolmogorov-Smirnov statistic). This pattern is evident when compared to a rank-ordered xAA background distribution of HOMO-LUMO gaps (Fig. 3B). The results demonstrated that C20 does not evenly disperse across this range. Instead, it is characterized by a large cluster of structurally stable, high E_g_ members (>11eV) with a targeted selection of high-gap outliers.

**Table 2.** HOMO-LUMO gap statistics.

| Metric | C20 | Random Alphabet Mean | Random Alphabet SD | C20 Percentile |
| --- | --- | --- | --- | --- |
| Range | 3.083 | 2.803 | 0.222 | 89 |
| Evenness | 0.059 | 0.032 | 0.014 | 5 |
| KS | 0.573 | 0.274 | 0.079 | 0 |

KS = Kolmogorov-Smirnov statistic; SD = standard deviation.

**Figure 3.**
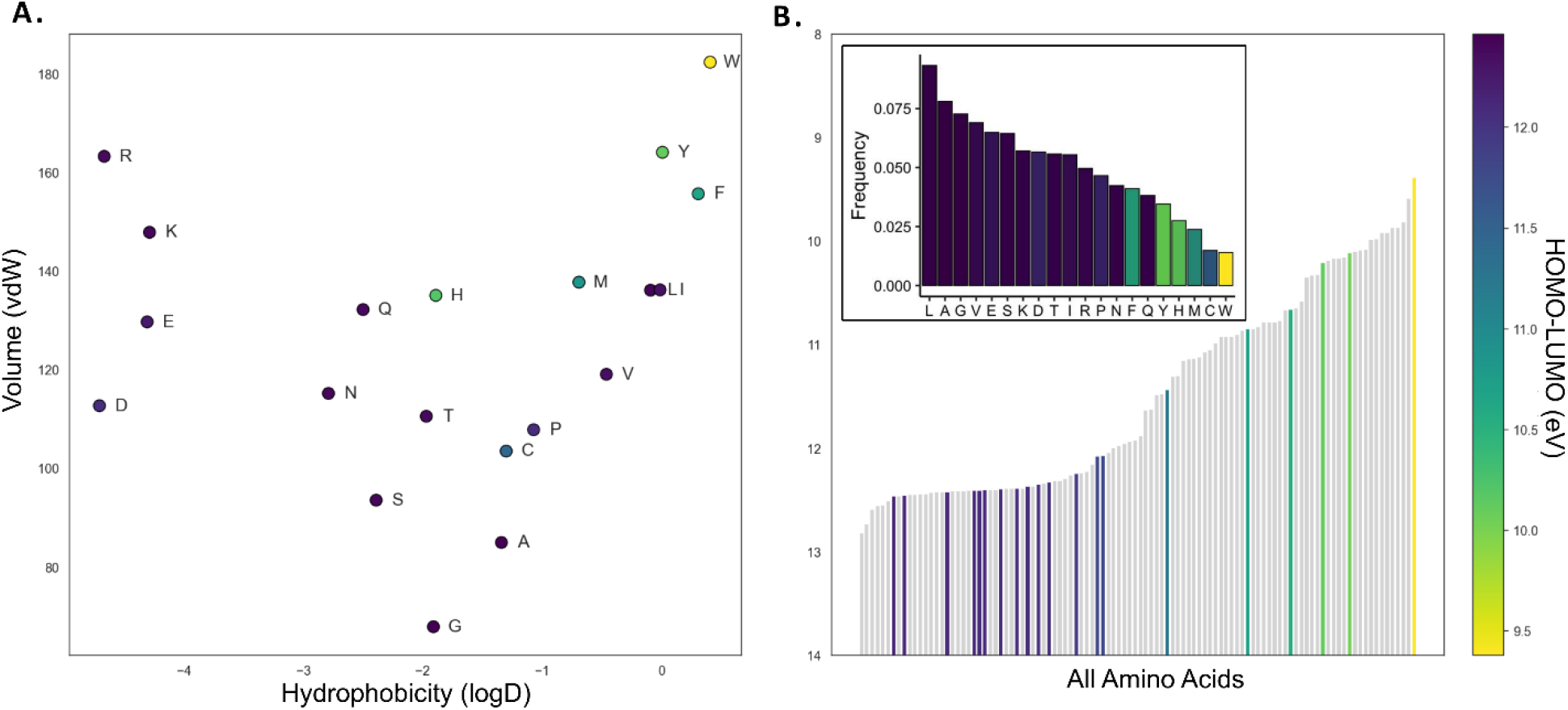
C20 in chemical space. **A.** C20 AAs in a three-dimensional chemical space – hydrophobicity measured by JChem LogD, van der Waals volume, and calculated HOMO-LUMO gaps (in color). **B**. Distribution of rank-ordered HOMO-LUMO gaps for C20 AAs (colored bars) relative to other xAAs (grey bars) and frequency bar plot of each canonical AA in every RCSB Protein Data Bank entry, colored by HOMO-LUMO gap (inset).

To contextualize the reactivity profile of C20 in greater detail, we isolated two alphabets from the background alphabet distribution: the sets with the minimum and maximum Earth Mover’s Distance (EMD or Wasserstein distance) to C20. Together, these two sets represented two extremes of sampled alphabets in terms of E_g_ (Fig. 4). The former successfully mimics the overall E_g_ distribution of C20 where constituents maintain both subtle similarity to and stark contrast from C20 (Fig. 4A-B). This set achieved a similar E_g_ profile to C20 by the incorporation of familiar functional groups (*e.g*. hydroxyls, amines, amides and aromatic rings) while simultaneously containing exotic side chains and heteroatoms (such as a nitril-, phosphate- and even adamantyl-bearing side chains). In other words, adherence to the particular distribution of HOMO-LUMO gaps found in C20 does not, by itself, prohibit significant chemical novelty. In contrast, the set with maximum EMD to C20 represents a statistically average, yet functionally distinct alphabet (Fig. 4A-B). Chemically, this set is quite distant to C20 with an overwhelming abundance of ring-bearing side chains, where 19 of 20 xAAs contained cyclic moieties (Fig. 4C). The abundance of cyclic side chains likely reflected the preponderance of ring-bearing side chains within the background library itself (which was indicative of successful random sampling of the background alphabets).

**Figure 4.**
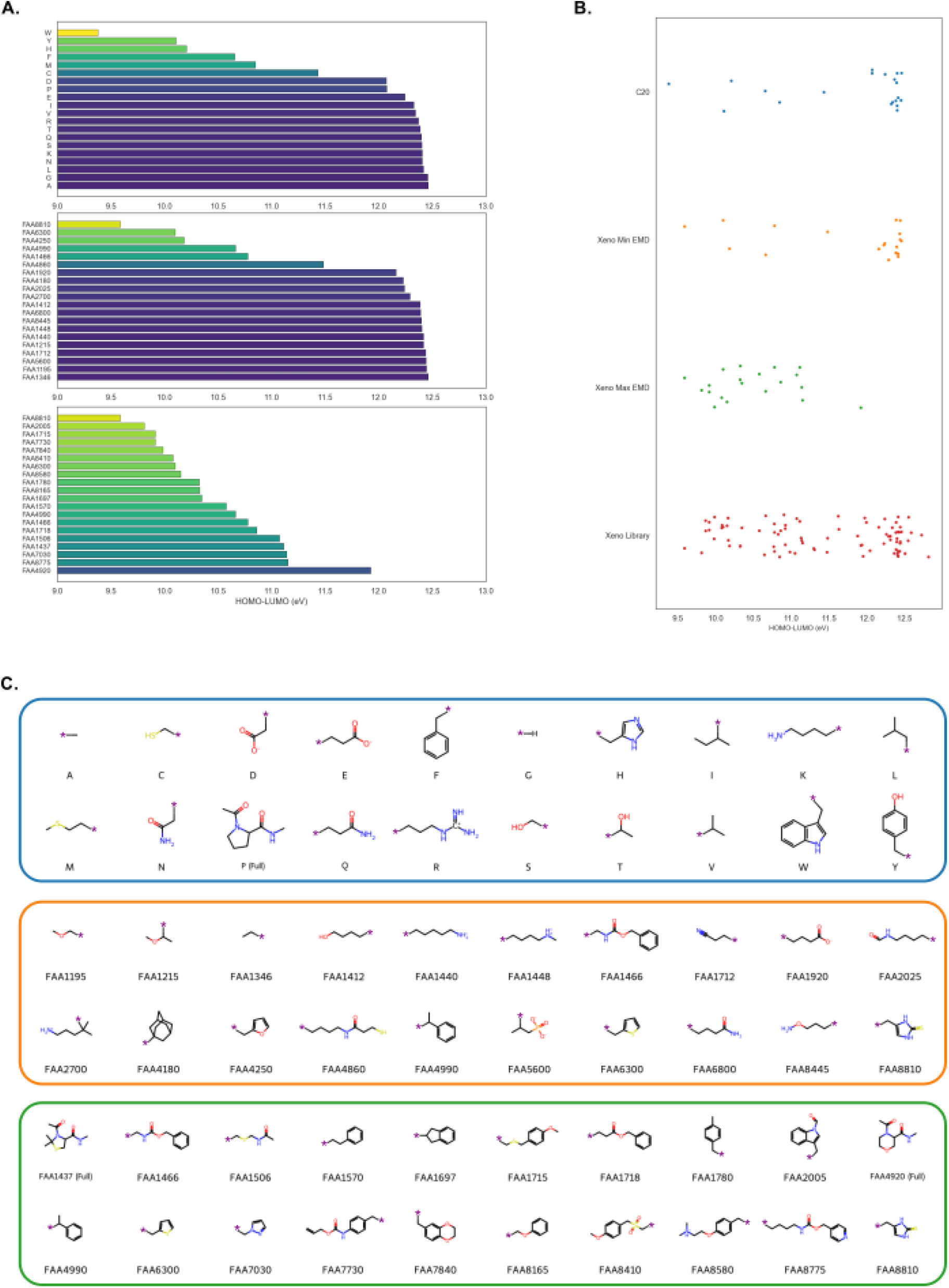
The HOMO-LUMO gap profile of C20 relative to xAA alphabets. **A.** Distribution of HOMO-LUMO gaps for C20 (top), a xeno alphabet with the lowest (middle) and highest (bottom) EMD to C20 from the library of 10 million random xeno alphabets. Bars colored by HOMO-LUMO gap values. **B**. Strip plots of HOMO-LUMO gaps for C20 (blue) and two xeno alphabets with the lowest (yellow) and highest (green) Earth Mover’s Distance (EMD) to C20 from the library of 10 million random xeno alphabets (red). **C**. Chemical structures of AA sidechains for C20 (top) and two xeno alphabets with the lowest (middle) and highest (bottom) EMD to C20. Purple asterisks denote Cα within the AA backbone, except for proline or proline-derivative amino acids where the whole amino acid is shown. One-letter codes for C20 or the IRIS Biotech (https://iris-biotech.de) Fmoc-L-α-monosubstituted amino acid catalogue number for xeno alphabets shown under each structure.

The core-and-outlier distribution of HOMO-LUMO gaps for C20 was consistent with the observed occurrence frequencies of AAs within both proteins as a whole and catalytic regions only (*i.e*. active sites).^18–20^ For instance, we measured the prevalence of AA residues in every protein within the Research Collaboratory for Structural Bioinformatics (RCSB) Protein Data Bank (PDB) (n=202,134 on 17 DEC 2025) and a clear trend between reactivity and frequency was observed (Pearson correlation = 0.73; Table 3, Fig. 3B): low E_g_ residues had significantly lower incorporation rates throughout an entire protein (Fig. 3B). When we examined which AAs were incorporated into the catalytic region(s) of proteins, an enrichment of low E_g_ residues was generally observed. Low E_g_ residues (*e.g*. W, H, Phenylalanine [F]), are found incorporated into the active site rather than dispersed at-random throughout the protein (Table 3). The results suggest that C20 may be particularly optimized for a core of high E_g_ residues with select low E_g_ residues, presumably for particular catalytic niches. Thus, C20 was not only optimized by natural selection for physicochemical properties related to structural propensity but also simultaneously constrained to a core-and-outlier reactivity distribution for specific catalytic functions.

**Table 3.** HOMO-LUMO gaps in enzyme active site residues.

| Enzyme Family | Aligned Residues (N) | Active Site Residues (N) | Mean Active Site HOMO-LUMO (eV) | Mean Non-Active Site HOMO-LUMO (eV) | P-Value |
| --- | --- | --- | --- | --- | --- |
| Carbonic anhydrase | 640 | 17 | 11.652 | 11.876 | 0.000019 |
| Cysteine protease papain | 543 | 5 | 11.732 | 11.899 | 0.63 |
| Heme peroxidase | 1,170 | 7 | 11.718 | 11.877 | 0.00006 |
| Lysozyme | 187 | 9 | 11.669 | 11.960 | 0.01 |
| RNase A | 1,718 | 11 | 11.620 | 11.796 | 0.0002 |
| Serine protease subtilisin | 505 | 3 | 11.561 | 12.045 | 0.000001 |
| Triosephosphate isomerase | 571 | 6 | 11.753 | 11.930 | 0.13 |

## Conclusions

The core-and-outlier distribution observed for C20 provides organisms with a unique inventory of stable structural building blocks while simultaneously permitting the incorporation of variably reactive side chains when compared to random alternatives. Natural selection likely optimized C20 for a functionally advantageous unevenness in terms of chemical reactivity. These results are consistent with observed AA incorporation in proteins and evidence surrounding the expansion of the genetic code as life evolved towards a single, universal set of building blocks.^3,17,21,22^ Our results align with over a decade of theoretical work suggesting that the distribution within chemistry space^23^ for C20 is distinctly non-random.^1–4,14^ We directly build upon this body of work by demonstrating that the non-random physicochemical distribution which was previously observed extends beyond hydrophobicity and volume into reactivity-relevant descriptors like HOMO-LUMO gaps. Together, these results reinforce the hypothesis that C20 is finely tuned for the construction of versatile and robust proteomes suited for nearly any environmental condition found on Earth.^24^

Interestingly, the results also demonstrate that sets of xAAs with similar functionally-relevant physicochemical patterns to C20 do indeed exist. One can imagine myriad applications for such xAA alphabets but three immediate use cases come to mind. First, this finding can serve to advance contemporary biotechnology and biomanufacturing by suggesting novel design strategies for the construction of (potentially) functional xeno proteins. While previous work^3,4^ has focused on constructing xeno peptides to evaluate the structure-forming potential of xeno alphabets, the field has yet to investigate *functional* alternatives to C20. This work specifically begins to inform researchers where the required constraints lie when attempting to design functional xeno proteins. It is plausible that the use of xAA alphabets containing novel sidechains and even heteroatoms could lead to the construction of proteins with emergent properties and the ability to catalyze reactions that are currently inaccessible. Second, the results introduce the HOMO-LUMO gap as a novel and potentially powerful feature for training machine learning models, including those that could be used to design xeno proteins. Third, this study may further astrobiology research by providing a guide on what to search for beyond C20. Instead of merely identifying the presence of certain canonical AAs, astrobiologists could instead search for *sets* of fundamental building blocks with unusual properties, *as a collective*, in terms of structure-forming and functional physicochemical properties.

While the model explored here demonstrates that xAA alphabets with physicochemical properties emulating C20 do exist, it will require a significant experimental leap to demonstrate that these alphabets can act within a functioning xenobiological system to maintain structural and functional biochemistry. A tractable next step would be to design and synthesize sets of 20 xAAs with this non-random profile so as to test these exact assumptions. Experiments here will be necessary to evaluate computational models and predictors, especially with the recent explosion of artificial intelligence in biotechnology. We anticipate that the results from these experiments would begin to answer two questions immediately: (i) Is the current model of physicochemical coverage sufficient to design functional xeno proteins? (ii) Do collections of xAAs able to form functional polymers exist?

## Methods

### Defining options for a library of xAAs

All analyses presented mined the same starting library of AAs used in a previous study.^4^ This collection of 248 monosubstituted L-α-amino acids was filtered down to 84 xAAs that were consistently returned in 10,000 heuristic searches for AA alphabets emulating the size and hydrophobicity profile of C20 (see Eq. 1). While previous work used a heuristic search for AA sets emulating a smaller primordial canonical alphabet,^4^ we refined this strategy to search for alphabets emulating the full set of genetically-encoded AAs (*i.e*. C20) with a similar heuristic fitness function:

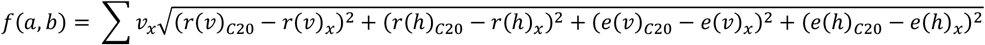

Where *x* is the xeno alphabet being evaluated, *v* is the calculated van der Waals volume, h is the calculated hydrophobicity (LogD, pH 7.0), e is the evenness of each descriptor (*v, h*), *r* is the range of a given descriptor, and Σ*v*_*X*_ is the sum of van der Waals volumes of all AAs in a given xeno alphabet. The weight metric Σ*v*_*X*_ ensures that alphabets mimic the specific range of volumes found within C20.

Molecular volume and logD (at pH 7.0) were calculated for each xAA (Table SI.1) and C20 AA (Table SI.2) using a previously described method.^3,25^ Briefly, JChem molecular volume was calculated for AAs with uncapped termini and hydrophobicity was calculated using JChem logD at pH 7.0 with the same AAs with an acetylated N-terminus and N-methylamidated C-terminus.

### Potential Energy Surfaces

To accurately model the behavior of AAs within proteins, we calculated the potential energy surface (PES) as previously described^4,26,27^ for each of these 84 xAAs, as well as C20, with acetylated N-termini and N-methylamidated C-termini (Fig. 1B-C). Each AA was considered in its native protonation state in water at pH 7.0. To generate initial conformers for every pair (N=144) of phi and psi dihedral angles (-180, -150, -120, -90, -60, -30, 0, 30, 60, 90, 120, 150; See Figure 1B and Supplementary Information), for each AA, random starting 3D geometries were generated in RDKit (v.2024.03.5, https://www.rdkit.org). Each was then optimized using the UFF force field^28^ with iteratively increasing force constraints (initial = 0.002, multiplier = 2.0 per iteration) for every desired phi and psi dihedral angles within a five-degree tolerance. UFF-optimized conformers were then further optimized in xTB at a GFN2-xTB level^29^ (--opt) with constraints on the dihedral angles (phi, psi, force constant = 0.05) using ALPB implicit solvation.^30^ The Conformer–Rotamer Ensemble Sampling Tool^31^ (CREST) program was then used to thoroughly sample every AA’s side chain for each conformation, while constraining backbone phi and psi angles to respective target values (force constant = 0.25). Finally, DFT single point energies were calculated in Orca^32^ (v.6.1.0) for every CREST conformer returned within 10 kcal⋅mol ^-1^, using the openCOSMO-RS,^33^ BP86 functional,^34^ def2-TZVP basis set,^35^ and Grimme’s D3(BJ) dispersion correlations.^36^ Final free energies of conformers (G) were calculated with the formula:

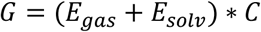

Where the final free energy *G* is equal to the sum of free energy of gas phase (*E*_*gas*_) and solvation (*E*_*solv*_) multiplied by the Hartree-to-kcal⋅mol^-1^ constant (*C*). The structure with the lowest DFT(BP86-D3BJ)/def2-TZVP//openCOSMO-RS energy was considered as the representative conformer for that phi and psi angle combination. The lowest DFT(BP86-D3BJ)/def2-TZVP//openCOSMO-RS energy across the entire PES for each AA was considered to be the relative minima for that AA and all other energies for that AA were scaled appropriately (Fig. 1C and Fig. SI.1).

### HOMO-LUMO gaps

We calculated the HOMO-LUMO gap energy for each AA where a PES was generated. Each conformer with the lowest DFT(BP86-D3BJ)/def2-TZVP//openCOSMO-RS energy within a PES was considered the representative conformer for that AA and thus underwent subsequent optimization without constraints at the DFT(BP86-D3BJ)/def2-TZVP//CPCMC(water)^37,38^ level in Orca. HOMO-LUMO gaps were then calculated from single-point energies on these optimized geometries at a HF^39–41^/def2-TZVPD//CPCM(water) level in Orca. These calculations were also performed with the same program versions and settings as presented above for the energy calculations.

### Analysis

All analyses were performed in Python (v3.11.11, https://www.python.org/). Statistical analyses and visualizations utilized scipy (v.1.15.3, https://scipy.org/), matplotlib (v.3.10.0, https://matplotlib.org/), numpy (v.2.2.5, https://numpy.org/), pandas (v.2.2.3, https://pandas.pydata.org/), and seaborn (v.0.13.2, https://seaborn.pydata.org/) Python libraries. The statistical non-randomness of C20 reactivity was evaluated through a Monte Carlo simulation (10,000,000 iterations), where random 20-member xeno alphabets were sampled from a dataset of 84 xAAs. For each alphabet, coverage was calculated as the range and evenness for each alphabet (see Fig. 1A and Brown *et al*.^4^ for how to calculate these values). The two-sided Kolmogorov-Smirnov test^42^ was additionally calculated as a proxy for uniformity using the scipy kstest function. Wasserstein distance^43^ (Earth Mover’s Distance - EMD) was additionally calculated as another measure for distributional similarity to C20 with the scipy wasserstein_distance function. EMD is a similarity metric used when comparing two distributions specifically by calculating the amount of ‘work’ required to move one distribution to match another.^44^ The percentile rank of C20 for each metric is then determined relative to the random background library distributions of the xAA alphabets.

All protein sequences within the RCSB PDB (www.rcsb.org) protein-only group (as of 17 DEC 2025, n=202,134) were downloaded, and the baseline frequency of each of the 20 canonical AAs was calculated to establish a background distribution. Subsequently, a curated set of nine enzyme families with active site annotations was analyzed to determine if residues with lower HOMO-LUMO gaps are enriched at these sites. For each family, sequences were compiled and subjected to multiple sequence alignment. Each AA in the alignments was then mapped to its corresponding HOMO-LUMO gap value. The Mann-Whitney U test was used to assess whether the distribution of these reactivity values at annotated active site positions was statistically different from that of all non-active site positions.

## Acknowledgements

S.M.B. acknowledges a postdoctoral fellowship through the National Research Council’s Research Associateship Program. The Department of Defense High Performance Computing Modernization Program (HPCMP) provided CPU hour allocations for the “High-End BioCompute for Biotechnology (HEBC4B)” program (Project 3265) to HPCMP PI J.H. This research was funded by the Office of Naval Research (WU# 61A1P4). The opinions and assertions contained herein are those of the authors and are not to be construed as those of the U.S. Navy, military service at large or U.S. Government. J.H., S.N.D., and G.J.V. are employees of the US Government.

## Author Contributions

Conceptualization: S.M.B., G.J.V.; Computational investigation: S.M.B., J.H., S.N.D.; Formal analysis: S.M.B.; Data curation: S.M.B., S.N.D.; Visualization: S.M.B., S.N.D.; Writing–original draft: S.M.B., S.N.D.; Writing–editing: S.M.B., J.H., S.N.D., G.J.V.; Supervision: S.N.D., G.J.V.; Project administration: S.N.D., G.J.V.; Funding acquisition: S.N.D., G.J.V.

## Data Availability Statement

All data needed to evaluate the conclusions in the paper are present in the paper and/or the Supplementary Information.

