## Supplementary Information for "The genetically-encoded amino acids distribute non-randomly within a functionally-relevant chemical space"

**Table SI.1.** Physicochemical descriptors of background xAAs.

| **IUPAC Chemical Name** | **SMILES** | **JChem Molecular Volume (Å^3^)** | **Capped JChem pH 7 LogD** | **HOMO-LUMO Gap (eV)** |
| --- | --- | --- | --- | --- |
| 2-amino-6-(1-carboxyethylamino)hexanoic acid | CC(NCCCCC(N)C(=O)O)C(=O)O | 210.13 | -3.47 | 12.819 |
| 2-amino-3-(dimethylamino)propanoic acid | CN(C)CC(N)C(=O)O | 132.12 | -2.84 | 12.728 |
| 2,3-diaminopropanoic acid | NCC(N)C(=O)O | 96.82 | -3.62 | 12.590 |
| 2,3-diaminobutanoic acid | CC(N)C(N)C(=O)O | 113.93 | -3.42 | 12.555 |
| 2-amino-3-(methylamino)propanoic acid | CNCC(N)C(=O)O | 114.38 | -3.88 | 12.549 |
| 2-amino-3-cyclohexyloxypropanoic acid | NC(COC1CCCCC1)C(=O)O | 184.87 | 0.05 | 12.508 |
| 2-aminobutanoic acid | CCC(N)C(=O)O | 101.88 | -0.82 | 12.460 |
| 2-amino-4-(methylamino)butanoic acid | CNCCC(N)C(=O)O | 131.28 | -4.86 | 12.445 |
| 2-amino-3-methoxypropanoic acid | COCC(N)C(=O)O | 111.14 | -1.75 | 12.444 |
| 2-amino-3-methyl-4-phosphonobutanoic acid | CC(CP(=O)(O)O)C(N)C(=O)O | 163.57 | -4.73 | 12.442 |
| 2-amino-4-cyanobutanoic acid | N#CCCC(N)C(=O)O | 118.68 | -1.64 | 12.438 |
| 2-aminohexanoic acid | CCCCC(N)C(=O)O | 135.85 | 0.07 | 12.428 |
| 2-amino-3-methoxybutanoic acid | COC(C)C(N)C(=O)O | 128.21 | -1.33 | 12.418 |
| 2,7-diaminoheptanoic acid | NCCCCCC(N)C(=O)O | 164.76 | -3.84 | 12.417 |
| 2-amino-4-methoxybutanoic acid | COCCC(N)C(=O)O | 128.1 | -1.69 | 12.417 |
| 2,5-diaminopentanoic acid | NCCCC(N)C(=O)O | 130.77 | -4.39 | 12.416 |
| 2-amino-5-methylhexanoic acid | CC(C)CCC(N)C(=O)O | 153.03 | 0.36 | 12.411 |
| 2-amino-6-aminooxyhexanoic acid | NOCCCCC(N)C(=O)O | 156.81 | -1.21 | 12.409 |
| 2-amino-6-(methylamino)hexanoic acid | CNCCCCC(N)C(=O)O | 165.31 | -4.11 | 12.401 |
| 2-amino-5-aminooxypentanoic acid | NOCCCC(N)C(=O)O | 139.91 | -1.66 | 12.398 |
| 2-amino-3-cyclopentylpropanoic acid | NC(CC1CCCC1)C(=O)O | 158.34 | 0.34 | 12.390 |
| 2,6-diamino-6-oxohexanoic acid | NC(=O)CCCC(N)C(=O)O | 149.25 | -2.06 | 12.390 |
| 2-amino-6-hydroxyhexanoic acid | NC(CCCCO)C(=O)O | 144.49 | -1.37 | 12.386 |
| 2-amino-6-(diaminomethylideneamino)hexanoic acid | N=C(N)NCCCCC(N)C(=O)O | 180.18 | -4.22 | 12.365 |
| 2-aminooctanedioic acid | NC(CCCCCC(=O)O)C(=O)O | 180.63 | -3.14 | 12.338 |
| 2-amino-4-methylsulfonylbutanoic acid | CS(=O)(=O)CCC(N)C(=O)O | 154.8 | -2.96 | 12.315 |
| 2-amino-2-cyclopentylacetic acid | NC(C(=O)O)C1CCCC1 | 141.44 | -0.03 | 12.314 |
| 2,6-diamino-3,3-dimethylhexanoic acid | CC(C)(CCCN)C(N)C(=O)O | 182.48 | -3.54 | 12.291 |
| 2-amino-2-cyclohexylacetic acid | NC(C(=O)O)C1CCCCC1 | 158.83 | 0.41 | 12.261 |
| 2-amino-6-formamidohexanoic acid | NC(CCCCNC=O)C(=O)O | 166.79 | -1.70 | 12.240 |
| 2-(1-adamantyl)-2-aminoacetic acid | NC(C(=O)O)C12CC3CC(CC(C3)C1)C2 | 205.22 | 0.73 | 12.227 |
| 2-aminohexanedioic acid | NC(CCCC(=O)O)C(=O)O | 146.65 | -3.79 | 12.160 |
| 2-amino-5-(cyclohexylamino)-5-oxopentanoic acid | NC(CCC(=O)NC1CCCCC1)C(=O)O | 223.39 | -0.48 | 12.044 |
| 2-amino-6-(prop-2-enoylamino)hexanoic acid | C=CC(=O)NCCCCC(N)C(=O)O | 193.19 | -0.90 | 11.996 |
| 2,5-diamino-3,3-dimethylpentanoic acid | CC(C)(CCN)C(N)C(=O)O | 165.53 | -3.63 | 11.974 |
| 3-amino-2,2-dimethylbutanedioic acid | CC(C)(C(=O)O)C(N)C(=O)O | 147.24 | -3.53 | 11.953 |
| 5,5-dimethylpyrrolidine-2-carboxylic acid | CC1(C)CCC(C(=O)O)N1 | 141.99 | -0.37 | 11.937 |
| morpholine-3-carboxylic acid | O=C(O)C1COCCN1 | 117.35 | -1.69 | 11.923 |
| piperidine-2-carboxylic acid | O=C(O)C1CCCCN1 | 125.14 | -0.63 | 11.881 |
| 2-amino-3-(prop-2-enoxycarbonylamino)propanoic acid | C=CCOC(=O)NCC(N)C(=O)O | 168.29 | -1.33 | 11.632 |
| 2-amino-5-(prop-2-enoxycarbonylamino)pentanoic acid | C=CCOC(=O)NCCCC(N)C(=O)O | 202.27 | -0.75 | 11.624 |
| 2-amino-6-(3-sulfanylpropanoylamino)hexanoic acid | NC(CCCCNC(=O)CCS)C(=O)O | 219.06 | -0.99 | 11.482 |
| 2-amino-5-[[amino-(hydroxyamino)methylidene]amino]pentanoic acid | N=C(NO)NCCCC(N)C(=O)O | 172.09 | -4.55 | 11.475 |
| 2-amino-6-(pyridine-3-carbonylamino)hexanoic acid | NC(CCCCNC(=O)c1cccnc1)C(=O)O | 233.32 | -1.02 | 11.309 |
| 2-amino-4-sulfanylbutanoic acid | NC(CCS)C(=O)O | 120.43 | -1.08 | 11.299 |
| 2-amino-6-(phenylmethoxycarbonylamino)hexanoic acid | NC(CCCCNC(=O)OCc1ccncc1)C(=O)O | 259.32 | -0.53 | 11.151 |
| 2-amino-3-pyrazol-1-ylpropanoic acid | NC(Cn1cccn1)C(=O)O | 135.38 | -1.47 | 11.141 |
| 2-amino-3-[(prop-2-enoxycarbonylamino)methylsulfanyl]propanoic acid | C=CCOC(=O)NCSCC(N)C(=O)O | 204.11 | -0.89 | 11.131 |
| 2,2-dimethyl-1,3-thiazolidine-4-carboxylic acid | CC1(C)NC(C(=O)O)CS1 | 144.03 | -0.58 | 11.114 |
| 3-(acetamidomethylsulfanyl)-2-aminopropanoic acid | CC(=O)NCSCC(N)C(=O)O | 168.37 | -2.24 | 11.073 |
| 2-amino-6-benzamidohexanoic acid | NC(CCCCNC(=O)c1ccccc1)C(=O)O | 237.42 | 0.20 | 11.050 |
| 2-amino-3-(2-carboxyethylsulfanyl)propanoic acid | NC(CSCCC(=O)O)C(=O)O | 165.46 | -3.90 | 10.983 |
| 4-(2-amino-2-carboxyethyl)sulfanylbutanoic acid | NC(CSCCCC(=O)O)C(=O)O | 182.44 | -3.88 | 10.924 |
| 2-amino-3-pyridin-3-ylpropanoic acid | NC(Cc1cccnc1)C(=O)O | 151.4 | -0.91 | 10.922 |
| 2-amino-3-(carboxymethylsulfanyl)propanoic acid | NC(CSCC(=O)O)C(=O)O | 148.47 | -4.54 | 10.921 |
| 2-amino-3-phenylmethoxypropanoic acid | NC(COCc1ccccc1)C(=O)O | 181.79 | -0.02 | 10.904 |
| 2-amino-5-oxo-5-phenylmethoxypentanoic acid | NC(CCC(=O)OCc1ccccc1)C(=O)O | 217.85 | 0.17 | 10.861 |
| thiomorpholine-3-carboxylic acid | O=C(O)C1CSCCN1 | 127 | -1.12 | 10.847 |
| 2-amino-4-(phenylmethoxycarbonylamino)butanoic acid | NC(CCNC(=O)OCc1ccccc1)C(=O)O | 229.57 | -0.27 | 10.828 |
| 2-amino-3-(phenylmethoxycarbonylamino)propanoic acid | NC(CNC(=O)OCc1ccccc1)C(=O)O | 212.52 | -0.33 | 10.779 |
| 2-amino-6-[bis(ethylamino)methylideneamino]hexanoic acid | CCN=C(NCC)NCCCCC(N)C(=O)O | 248.71 | -3.01 | 10.778 |
| 2-amino-4-phenylmethoxybutanoic acid | NC(CCOCc1ccccc1)C(=O)O | 198.81 | 0.04 | 10.777 |
| 2-amino-4-butylsulfanylbutanoic acid | CCCCSCCC(N)C(=O)O | 188.64 | 0.53 | 10.769 |
| 2-amino-3-phenylbutanoic acid | CC(c1ccccc1)C(N)C(=O)O | 172.87 | 0.68 | 10.662 |
| 1,2,3,4-tetrahydroisoquinoline-3-carboxylic acid | O=C(O)C1Cc2ccccc2CN1 | 161.74 | 0.29 | 10.646 |
| 2-amino-4-phenylbutanoic acid | NC(CCc1ccccc1)C(=O)O | 172.6 | 0.76 | 10.578 |
| 2-amino-2-(2,3-dihydro-1H-inden-2-yl)acetic acid | NC(C(=O)O)C1Cc2ccccc2C1 | 177.85 | 0.73 | 10.347 |
| 2-amino-3-phenoxypropanoic acid | NC(COc1ccccc1)C(=O)O | 164.75 | -0.06 | 10.327 |
| 2-amino-3-(4-methylphenyl)propanoic acid | Cc1ccc(CC(N)C(=O)O)cc1 | 172.49 | 0.83 | 10.324 |
| 2-amino-3-(furan-2-yl)propanoic acid | NC(Cc1ccco1)C(=O)O | 136.8 | -0.86 | 10.183 |
| 2-amino-3-[(4-methylphenyl)methylsulfanyl]propanoic acid | Cc1ccc(CSCC(N)C(=O)O)cc1 | 208.2 | 1.23 | 10.168 |
| 2-amino-3-(3-methoxyphenyl)propanoic acid | COc1cccc(CC(N)C(=O)O)c1 | 181.68 | 0.16 | 10.164 |
| 2-amino-3-[4-[2-(dimethylamino)ethoxy]phenyl]propanoic acid | CN(C)CCOc1ccc(CC(N)C(=O)O)cc1 | 245.88 | -1.60 | 10.150 |
| 2-amino-3-thiophen-2-ylpropanoic acid | NC(Cc1cccs1)C(=O)O | 146.94 | 0.23 | 10.098 |
| 2,3-dihydro-1H-indole-2-carboxylic acid | OC(=O)C1Cc2ccccc2N1 | 143.25 | 0.22 | 10.085 |
| 2-amino-3-[(4-methoxyphenyl)methylsulfonyl]propanoic acid | COc1ccc(CS(=O)(=O)CC(N)C(=O)O)cc1 | 234.51 | -1.30 | 10.080 |
| 2-amino-3-(2,3-dihydro-1,4-benzodioxin-6-yl)propanoic acid | NC(Cc1ccc2c(c1)OCCO2)C(=O)O | 196.61 | -0.17 | 9.985 |
| 2-amino-3-[4-(carboxymethoxy)phenyl]propanoic acid | NC(Cc1ccc(OCC(=O)O)cc1)C(=O)O | 209.33 | -3.49 | 9.979 |
| 2-amino-3-[4-(prop-2-enoxycarbonylamino)phenyl]propanoic acid | C=CCOC(=O)Nc1ccc(CC(N)C(=O)O)cc1 | 238.93 | 0.90 | 9.915 |
| 2-amino-3-[(4-methoxyphenyl)methylsulfanyl]propanoic acid | COc1ccc(CSCC(N)C(=O)O)cc1 | 217.34 | 0.56 | 9.915 |
| 2-amino-3-phenylsulfanylpropanoic acid | NC(CSc1ccccc1)C(=O)O | 174.47 | 0.47 | 9.868 |
| 2-amino-3-(4-methoxy-2,6-dimethylphenyl)propanoic acid | COc1cc(C)c(CC(N)C(=O)O)c(C)c1 | 215.81 | 1.18 | 9.866 |
| 2-amino-3-(1-formylindol-3-yl)propanoic acid | NC(Cc1cn(C=O)c2ccccc12)C(=O)O | 202.21 | -0.32 | 9.813 |
| 2-amino-3-(2-sulfanylidene-1,3-dihydroimidazol-4-yl)propanoic acid | NC(Cc1c[nH]c(=S)[nH]1)C(=O)O | 152.83 | -1.44 | 9.588 |

**Table SI.2.** C20 physicochemical descriptors.

| **Amino Acid** | **1-Letter Code** | **SMILES** | **JChem**  **Molecular Volume (Å^3^)** | **Capped**  **JChem**  **pH 7 LogD** | **HOMO-LUMO**  **Gap (eV)** |
| --- | --- | --- | --- | --- | --- |
| G | G | NCC(=O)O | 67.85 | -1.91 | 12.4557 |
| T | T | CC(O)C(N)C(=O)O | 110.52 | -1.97 | 12.3861 |
| Q | Q | NC(=O)CCC(N)C(=O)O | 132.15 | -2.5 | 12.396 |
| I | I | CCC(C)C(N)C(=O)O | 136.14 | -0.01 | 12.3256 |
| R | R | N=C(N)NCCCC(N)C(=O)O | 163.21 | -4.67 | 12.3696 |
| D | D | NC(CC(=O)O)C(=O)O | 112.66 | -4.71 | 12.0715 |
| M | M | CSCCC(N)C(=O)O | 137.7 | -0.69 | 10.8474 |
| Y | Y | NC(Cc1ccc(O)cc1)C(=O)O | 164.03 | 0.01 | 10.1096 |
| C | C | NC(CS)C(=O)O | 103.46 | -1.3 | 11.4336 |
| S | S | NC(CO)C(=O)O | 93.51 | -2.39 | 12.4032 |
| K | K | NCCCCC(N)C(=O)O | 147.81 | -4.29 | 12.4037 |
| F | F | NC(Cc1ccccc1)C(=O)O | 155.62 | 0.31 | 10.6567 |
| L | L | CC(C)CC(N)C(=O)O | 136.08 | -0.09 | 12.4171 |
| V | V | CC(C)C(N)C(=O)O | 119.03 | -0.46 | 12.343 |
| P | P | C1CC(NC1)C(=O)O | 107.79 | -1.07 | 12.0731 |
| E | E | NC(CCC(O)=O)C(O)=O | 129.65 | -4.31 | 12.2442 |
| W | W | NC(Cc1c[nH]c2ccccc12)C(O)=O | 182.3 | 0.41 | 9.3793 |
| A | A | CC(N)C(O)=O | 84.93 | -1.34 | 12.4623 |
| N | N | NC(CC(N)=O)C(O)=O | 115.16 | -2.79 | 12.404 |
| H | H | NC(CC1=CNC=N1)C(O)=O | 135.02 | -1.89 | 10.2074 |

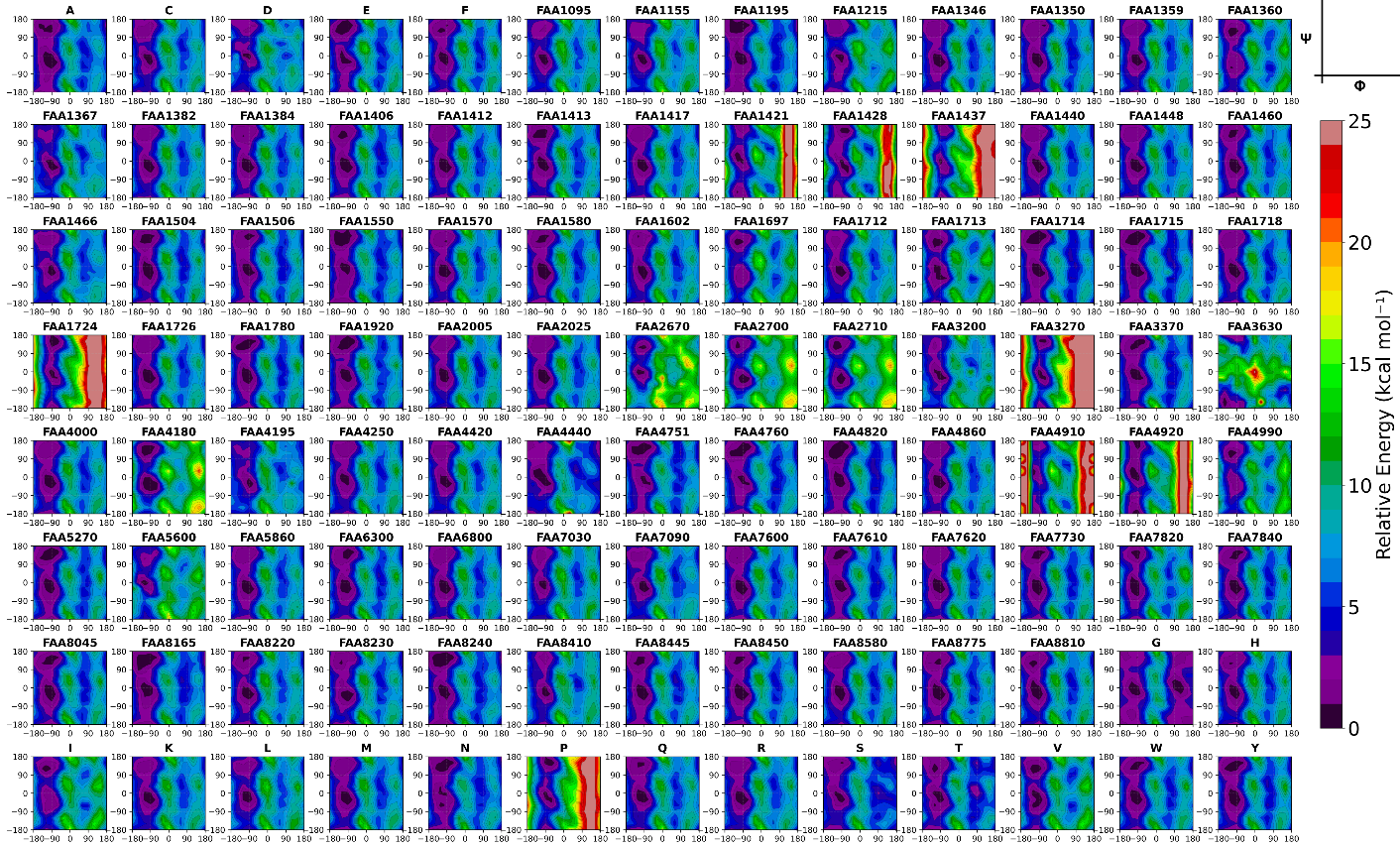

**Figure SI.1** Potential Energy Surfaces for all AAs calculated. Energies shown on a linear color map from black (0 kcal/mol) to red (25 kcal/mol).
