## Supplementary material for "The genetically-encoded amino acids distribute non-randomly within a functionally-relevant chemical space": PES_SI.html

Interactive Scatter Plots


Distribution Statement A. Approved for public release: distribution is unlimited.

### Ramachandran Scatter Plots

Distribution Statement A. Approved for public release: distribution is unlimited.
